# Investigation of novel functions of three genes in oriental river prawn, *Macrobrachium nipponense*: Molecular Cloning, Expression, and *In situ* Hybridization Analysis

**DOI:** 10.1101/441147

**Authors:** Shubo Jin, Hongtuo Fu, Yuning Hu, Shengming Sun, Sufei Jiang, Yiwei Xiong, Hui Qiao, Wenyi Zhang, Yongsheng Gong, Yan Wu

## Abstract

Three genes were predicted to be potentially involved in the male sexual development in *M*. *nipponense*, including the Gem-associated protein 2-like isoform X1 (GEM), Ferritin peptide, and DNA polymerase zeta catalytic subunit (Rev3). In this study, we aimed to investigate their novel functions in depth. The full-length cDNA sequence of Mn-GEM was 1,018 bp, encoding 258 amino acids. The partial Mn-Rev3 cDNA sequence was 6,832 bp, encoding 1,203 amino acids. Tissue distribution indicated that all of these three genes have higher expression level in testis and androgenic gland, implying their novel functions in male sexual development. In situ hybridization analysis further confirmed the novel roles of these three genes. Rev3 promote the testis development during the whole reproductive cycle, while GEM and ferritin only promote the activation of testis development. Besides, these three genes play essential roles in funicular structure development surrounding the androgenic gland cells, which promote and support the formation of androgenic gland cells. The expression in hepatopancreas cells also suggested their role in immune system in *M. nipponense.* This study advances our understanding of male sexual development in *M. nipponense*, as well as providing the basis for further studies of male sexual differentiation and development in crustaceans.

## Introduction

The oriental river prawn, *Macrobrachium nipponense* (Crustacea; Decapoda; Palaemonidae), are widely distributed in freshwater and low-salinity estuarine regions of China and other Asian countries [1–5]. It is a commercially important species with an annual aquaculture production of 205,010 tons [6]. The growth performance of male *M. nipponense* and female *M. nipponense* showed significant difference during the *M. nipponense* aquaculture. “Male prawns of *M. nipponense* grow faster and reach larger size at the harvest time than their female counterparts” [4–5]. The establishment of an artificial techniques to produce all male progeny on a commercial scale is a long-term goal in *M. nipponense* aquaculture. Therefore, it is urgently needed to fully understand the sex differentiation and determination mechanisms of *M. nipponense*.

The androgenic gland is found in most crustaceans, producing hormones, which promote the driving of male sexual differentiation, the establishment of male sexual characteristics, and the development of the testes [7]. The ablation of androgenic gland from male *Macrobrachium rosenbergii* prawn results in the sex reversal to the “neo-females”. All male progeny was generated when the “neo-females” were mated with normal male *M. rosenbergii* [7–9]. Thus, studies on androgenic gland is a hot topic on male sexual differentiation and development in crustacean species. The genes in the androgenic gland is of great importance to be fully understood, especially for those which may promote the male differentiation and development. The androgenic gland transcriptome and miRNA library have both been constructed for *M. nipponense* [10–11]. A series of genes identified in the androgenic gland transcriptome have been analyzed and proven to be involved in the sex-differentiation and determination mechanism ofM. *nipponense* [12–15].

iTRAQ technique was used to perform the quantitative proteomic analysis of androgenic gland during the nonreproductive and reproductive season in *M. nipponense.* A total of 3 differentially expressed proteins (DEPs) showed highest expression level in androgenic gland, compared with those in testis and ovary, including Gem-associated protein 2-like isoform X1 (GEM), Ferritin peptide, and DNA polymerase zeta catalytic subunit (Rev3) [16]. According to the previous studies, ferritin peptide and Rev3 have been proven to be involved in the immune system maintenance. Ferritin peptide has been analysed in *M. nipponense*, which plays essential roles in its innate immune defence, especially for that in cellular and organismic iron homeostasis [17]. However, these 3 DEPs might play essential roles in male sexual development in *M. nipponense*, based on their higher expression levels in androgenic gland and the importance of the androgenic gland in male sex differentiation and sex determination in crustacean species. The roles in male sexual development might be the novel functions that did not report yet.

In this study, we aimed to further analyse their functions in *M. nipponense* in depth, especially for the important roles in male sexual roles. The full-length cDNA sequences from *M. nipponense* were cloned and their structural characteristics were analysed. The mRNA expression patterns in different tissues and reproductive cycle of testis were determined by quantitative real-time PCR (qPCR), and their locations were further determined by *in situ* hybridization. The results of this study provide the foundations for male sexual development in *M. nipponense*, as well as that in other crustacean species.

## Materials and methods

### Ethics Statement

As described in detail previously [10], the prawns were obtained from the Tai Lake in Wuxi, China. We got the permission from the Tai Lake Fishery Management Council. *M. nipponense* is a normal species with huge production in China, which can be used for experimental materials. All the experimental animal programs involved in this study were followed the experimental basic principles, approved by committee of Freshwater Fisheries Research Institute. MS222 anesthesia was used for each prawn when androgenic glands were collected, in order to minimize suffering.

### Prawn and Tissue Preparation

As described in detail previously [13], healthy adult *M. nipponense* with wet weight of 3.785-26g were obtained from Tai Lake in Wuxi, China (120°13’44“E, 31°28’22”N). These specimens were maintained in aerated freshwater under lab conditions at the temperature of 28°C for at least 72 h prior to tissue collection. A total of 6 tissues were collected from mature prawns for qPCR analysis, including ovaries, testes, androgenic glands, heart, intestine and hepatopancreas, in order to determine the mRNA expression levels in different tissues. An additional androgenic gland was collected for Rapid Amplification of cDNA Ends (RACE) cloning. The Olympus SZX16 dissecting microscope was used to extract the androgenic glands. Testis in reproductive season at the temperature of 28°C and testis in non-reproductive season at the temperature of 15°C were collected, in order to determine the expression levels in reproductive cycle of testis. The samples were treated with phosphate buffer saline (PBS), and immediately frozen in liquid nitrogen until used for RNA extraction to prevent total RNA degradation.

### Rapid Amplification of cDNA Ends (RACE)

As described in detail previously [13], total RNA was extracted from androgenic gland as template using RNAiso Plus Reagent (Takara Bio Inc.), followed the protocol of the manufacturer. The RNase-free DNase I (Sangon, Shanghai, China) was used to treat the isolated RNA to eliminate possible genomic DNA contamination. BioPhotometer (Eppendorf, Hamburg, Germany) was used to measure the concentration of the total RNA sample with the A260/A280 in the range of 1.8-2.0. The RNA quality was then measured by 1% agarose gel.

As described in detail previously [13], a M-MLV reverse transcriptase was used to perform the first strand 3′cDNA and 5′cDNA synthesis for gene cloning using the 3′-Full RACE Core Set Ver.2.0 kit and the 5′-Full RACE kit (Takara Bio Inc., Japan), respectively with the reaction conditions recommended by the manufacturer. The synthesized cDNAs were kept at -20°C. 3′/5′-RACE PCR reactions were performed with the 3′ gene-specific primer (GEM-3GSP1, GEM-3GSP2, Rev3-3GSP1, Rev3-3GSP2) or 5′GSP (GEM-5GSP1, GEM-5GSP2) (Table.1). The partial unigene sequences were obtained from the *M. nipponense* androgenic gland transcriptome, and the 3′GSP and 5′GSP of each gene were designed based on the unigene sequence. 1% agarose gel was used to measure the PCR product.

As described in detail previously [13], the Gel Extraction kit (Sangon, shanghai,China) was used to cut and purify the PCR products, following the manufacturer’s instructions. Amplified cDNA fragments were transferred into the pMD18-T vector (Takara Bio Inc., Japan). Recombinant bacteria were identified by blue/white screening and confirmed by PCR. An automated DNA sequencer (ABI Biosystem, USA) was used to determine the nucleotide sequences of the cloned cDNAs. BLAST software (http://www.ncbi.nlm.nih.gov/BLAST/) was used to examine the nucleotide sequence similarities.

### Nucleotide Sequence and Bioinformatics Analyses

As described in detail previously [13], the primer designing tool (http://www.ncbi.nlm.nih.gov/tools/primer-blast/) was used to design all primers used in this experiment. The 5′ and 3′ sequences from RACEs were assembled with the partial cDNA sequence corresponding to each fragmental sequence by DNAMAN 5.0. The BLASTX and BLASTN search program (http://www.ncbi.nlm.nih.gov/BLAST/) of GenBank was used to analyse the sequences based on the nucleotide and protein databases using. The ORF Finder tool (http://www.ncbi.nlm.nih.gov/gorf/gorf.html) was used to predict the open reading frame. ClustalW1.81 was used to perform multiple sequence alignment. Molecular Evolutionary Genetics Analysis, MEGA 5.1 was used to construct the phylogenetic trees based on the amino acid sequences using the neighbor-joining method.

### Various Reproductive Cycle of Testis and Various Tissues Expression by qPCR

As described in detail previously [13], qPCR was used to measure the relative mRNA expression of Mn-GEM, Mn-Rev3 and Mn-Ferritin at various adult tissues and various reproductive cycle of testis. Each tissue sample was dissected out from at least 3 mature prawns. RNAiso Plus Reagent (TaKaRa) was used to exact and isolate the total RNA from various tissues of adult prawns and different reproductive cycle of testis, following the manufacturer’s instructions. The concentration and quality of each total RNA sample was measured by BioPhotometer (Eppendorf) with A260/A280 as 1.8-2.0, and 1% agarose gel. Experiments were performed in triplicate. Approximately 1μg of total RNA from each tissue was used for the first-strand cDNA synthesis using iScriptTM cDNA Synthesis Kit perfect Real Time (Bio-Rad, CA, USA) and following the manufacturer’s instructions. The synthesized cDNA template for qPCR was kept at −20°C. The qPCR primers of each gene were designed based on the open reading frame. The Bio-Rad iCycler iQ5 Real-Time PCR System (Bio-Rad) was used to carry out the SYBR Green RT-qPCR assay. β-actin was used as an internal reference to amplify for the same sample (the primer’s sequences are shown in Table 1). Diethypyrocarbonate-water (DEPC-water) for the replacement of template was used as a negative control. All samples were run in triplicate (each duplicate for target gene and β-actin gene). The relative mRNA expression of each gene was calculated based on the 2^−⊿⊿CT^ comparative CT method. There are many advantages of the 2^−⊿⊿CT^ comparative CT method, including the ease of use and the ability of the data as “fold change” in expression. The amplification efficiency of target gene and β-actin were estimated by qPCR, using different concentrations of androgenic gland template. The androgenic gland templates include undiluted, two times diluted, four times diluted and eight times diluted sample. The slope of the Mn-GEM and β-actin at different concentrations of diluted samples were 1.393 and 1.411, respectively. The slope of the Mn-Rev3 and β-actin at different concentrations of diluted samples were 1.479 and 1.458, respectively. The slope of the Mn-Ferritin and β-actin at different concentrations of diluted samples were 1.576 and 1.583, respectively. Thus, the amplification efficiency between the target gene and β-actin are the same in this study. The tissue with lowest expression level was setted as 1 (a relative criterion), and other tissues were then compared with the relative criterion.

**Table 1.**
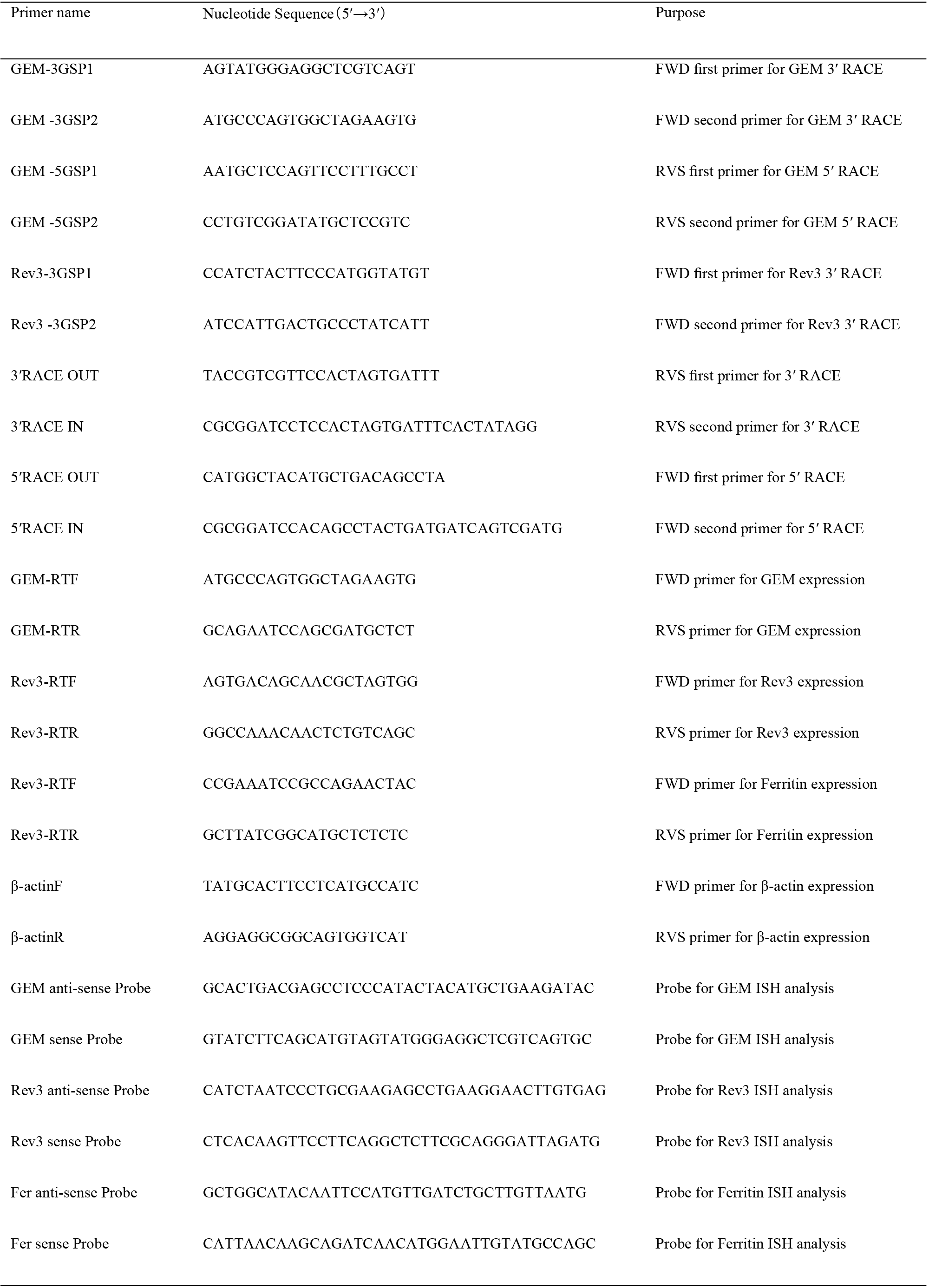
Universal and specific primers used in this study

### In situ Hybridization of DEM, Rev3 and Ferritin peptide mRNA in testis, androgenic gland and hepatopancreas

As described in detail previously [13], 4% paraformaldehyde treated by DEPC water was used to fix the tissue samples for *in situ* hybridization study. Primer5 software was used to design the anti-sense and sense probes of CISH (Chromogenic in-situ hybridization) study with DIG signal based on the cDNA sequence of each gene. The anti-sense and sense probes were then synthesized by Shanghai Sangon Biotech Company. Both of the anti-sense and sense probes were hybridized with the slide. The anti-sense probe and sense probe were prepared for the experimental group and control group, respectively. CISH study was performed on 4μ thick formalin fixed paraffin-embedded sections using Zytofast PLUS CISH implementation kit (Zyto Vision GmBH, Germany). As described in detail previously [13], a standard deparaffinization technique was performed by 10 min incubation in 3% H_2_O_2_. Following rinsing in deionized water (DW), target retrieval was achieved using pepsin digestion in a humidity chamber for 10 min. Slides were incubated in EDTA solution at 95 for 15 min after washing in DW. Slides were washed in DW and drained off; 20 μl of CISH anti-sense probe and sense probe were poured over each slide. Denaturation at 75 for 5 min was subsequently followed by hybridization at 37 for 60 min in the Thermobrite TM hybridization chamber (Vysis Inc., USA). Tris-buffered-saline (TBS) washing, at 55 and room temperature, each for five min was done concurrently. Mouse-anti-DIG (Zyto Vision GmBH, Germany) was poured drop-wise over each slide, and incubated in a humidity chamber at 37 for 30 min. Three washings, each for a minute with TBS was done, before and after incubating slides in anti-mouse-HRP-polymer for 30 min at room temperature. 3,3’-diaminobenzidine (DAB) solution was prepared as per guidelines (Zytofast PLUS CISH) and poured 50ul in each slide for 10 min at room temperature. After washing, a hematoxylin was used for counterstaining. Slides were dehydrated in graded alcohol solutions, air dried and mounted with mixture of distyrene, plasticizer and xylene (DPX). Slides were examined under light microscope for evaluation.

### Statistical Analysis

Quantitative data were expressed as mean ± SD. Statistical differences were estimated by oneway ANOVA followed by LSD and Duncan’s multiple range test. All statistics were measured using SPSS Statistics 13.0. A probability level of 0.05 was used to indicate significance (*P* < 0.05).

## Results

### Sequences analysis

The full-length Mn-DEM cDNA sequence was 1,018 bp with an open reading frame of 777 bp, encoding 258 amino acids. The 5′ and 3′ untranslated regions (UTRs) of Mn-DEM contained 147 bp and 94 bp, respectively. The partial Mn-Rev3 cDNA sequence was 6,832 bp with an open reading frame of 3,612 bp, encoding 1,203 amino acids. The 3′ untranslated regions (UTRs) of Mn-Rev3 contained 3220. The cDNA sequence of Mn-DEM and Mn-Rev3 have been submitted to GenBank with the accession no.MH817847 and MH817848, respectively. The relative information for Mn-Ferritin can be seen in previous study (Sun et al., 2014).

### Similarity comparison and phylogenetic analysis

The species used for GEM amino acid sequence blast have been listed in Table 2. The identities between Mn-GEM and GEM in other species was 42%-47% revealed by the BLASTP similarity comparisons, while the query coverage reached to 98%. Mn-GEM has the highest identity with GEM sequence from *Acanthaster planci.* MEGA 5.1 was used to construct a condensed phylogenetic tree using the neighbour-joining method, in order to analyse the evolutionary relationship between Mn-GEM and other well-defined GEM sequences, based on their completed amino acid sequences deposited in NCBI. The phylogenetic tree generated two main branches; one including amino acid sequences from different species, and another separate branch only including *M. nipponense* (Figure 1-A). Mn-GEM has dramatically long evolutionary relationship with those from other species.

**Table 2.** Amino acid sequence used for phylogenetic analysis of GEM

| Species | Accession Number |
| --- | --- |
| *Macrobrachium nipponense* |  |
| *Acanthaster planci* | XP_022103928.1 |
| *Xenopus laevis* | XP_018087535.1 |
| *Stylophora pistillata* | XP_022804116.1 |
| *Odobenus rosmarus divergens* | XP_004399834.1 |
| *Pomacea canaliculate* | XP_025106566.1 |
| *Centruroides sculpturatus* | XP_023219163.1 |
| *Biomphalaria glabrata* | XP_013071285.1 |
| *Aplysia californica* | XP_005105416.1 |
| *Mizuhopecten yessoensis* | XP_021367489.1 |
| *Xenopus tropicalis* | NP_001096228.1 |
| *Xenopus laevis* | NP_001087945.2 |
| *Mus musculus* | NP_079932.2 |
| *Mus caroli* | XP_021034811.1 |

**Fig 1:**
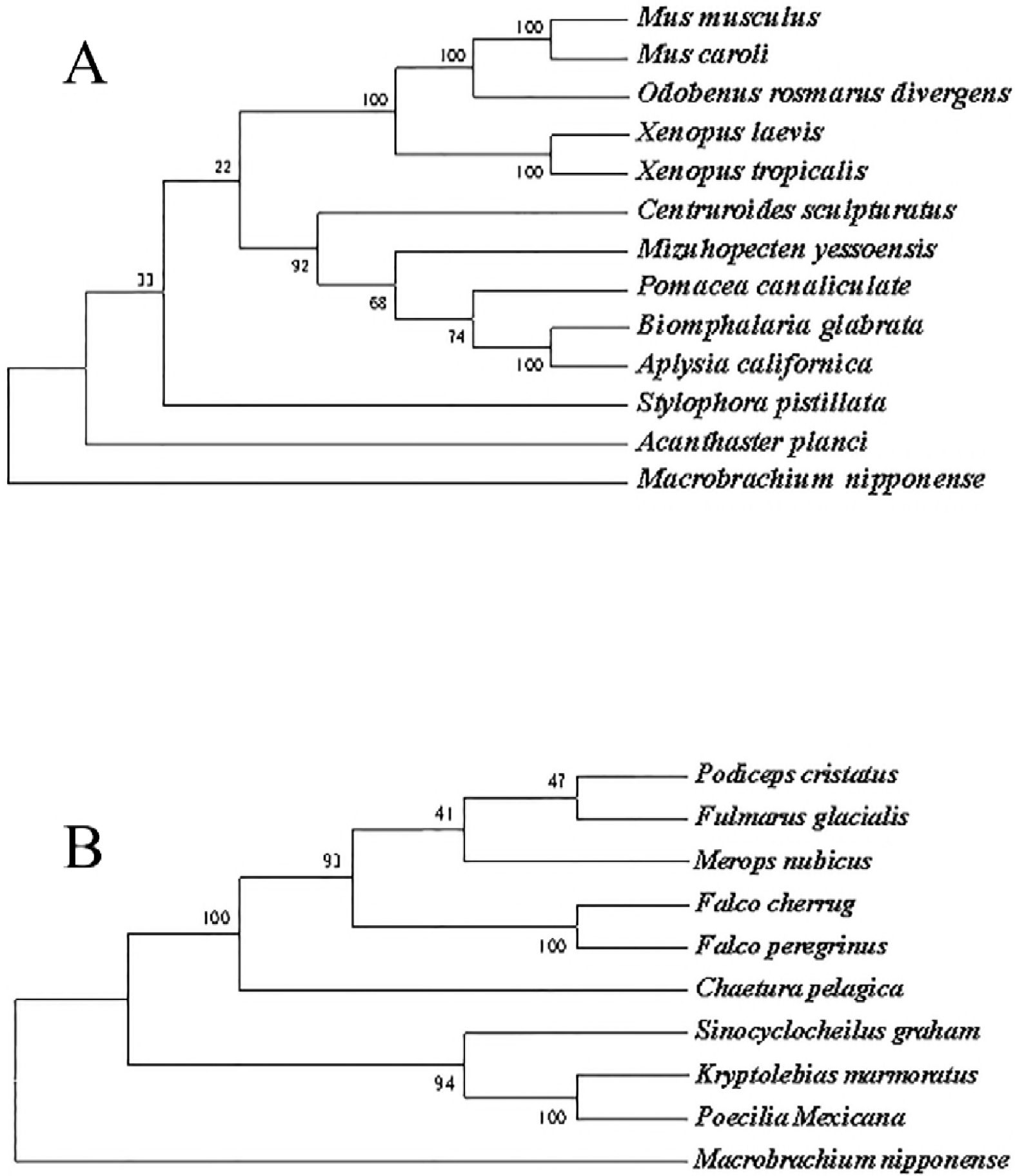
The construction of phylogenetic tree from different organisms based on amino acid sequence comparisons. Species names are listed on the right of the tree. A indicated the phylogenetic tree of GEM; B indicated the phylogenetic tree of Rev3.

The species used for Rev3 amino acid sequence blast have been listed in Table 3. The identities between the Mn-Rev3 and Rev3 in other species was 39%-41%, and the query coverage is only 35%. Mn-Rev3 has the highest identity with Rev3 sequence from *Orbicella faveolata.* Similar with that of Mn-GEM, *M. nipponense* is in a separate branch (Figure 1-B), revealed by phylogenetic tree using MEGA 5.1. Mn-GEM has dramatically long evolutionary relationship with those from other species.

**Table 3.** Amino acid sequence used for phylogenetic analysis of GEM

| Species | Accession Number |
| --- | --- |
| *Macrobrachium nipponense* |  |
| *Kryptolebias marmoratus* | XP_024866820.1 |
| *Chaetura pelagica* | KFU85727.1 |
| *Falco cherrug* | XP_014140871.1 |
| *Podiceps cristatus* | KFZ50056.1 |
| *Merops nubicus* | KFQ35536.1 |
| *Fulmarus glacialis* | XP_009579679.1 |
| *Sinocyclocheilus graham* | XP_016087315.1 |
| *Poecilia Mexicana* | XP_014866147.1 |
| *Falco peregrinus* | XP_013160015.1 |

### Expression analysis in different tissues and reproductive cycle of testis

Tissue distribution may reflect the physiological function of a protein. qPCR was used to determine the tissue distributions of these three genes. According to the qPCR analysis, the mRNA expression of these three genes in testis and androgenic gland were higher than that in ovary (Figure 2). The expressions showed significant difference for Rev3 and ferritin peptide (p<0.05). Besides, the mRNA expression of Rev3 and ferritin showed high expression level in hepatopancreas, especially for that of Rev3. The expression was 22.1-fold higher than that in heart (p<0.01).

**Fig 2:**
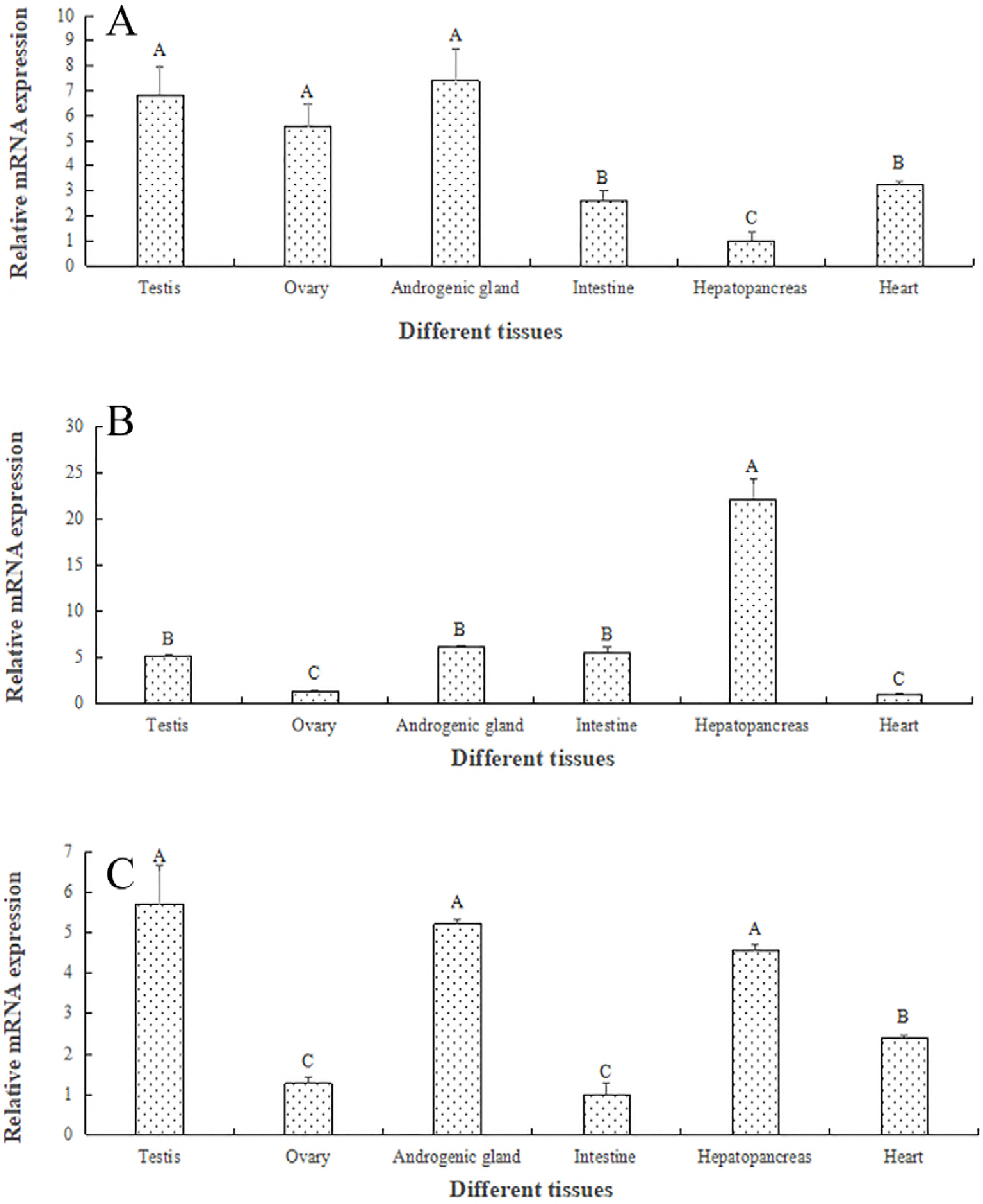
Expression characterizations in the various adult tissues were revealed by real-time quantitative PCR. The amount of mRNA was normalized to the β-actin transcript level. Data are shown as mean ±SD (standard deviation) of tissues in three separate individuals. Capital letters indicate expression difference of <u>Mn-Foxl2</u> in different adult tissues. A indicated the expression characterization of GEM; B indicated the expression characterization of Rev3; C indicated the expression characterization of ferritin peptide.

The mRNA expression of these three genes showed higher expression levels in reproductive season of testis than those in non-reproductive season and showed significant difference (p<0.05). The mRNA expressions of Mn-GEM, Mn-Rev3, and Mn-Ferritin in reproductive season were 2.31-fold, 2.97-fold and 2.67-fold higher than those in nonreproductive season (Figure 3).

**Fig 3:**
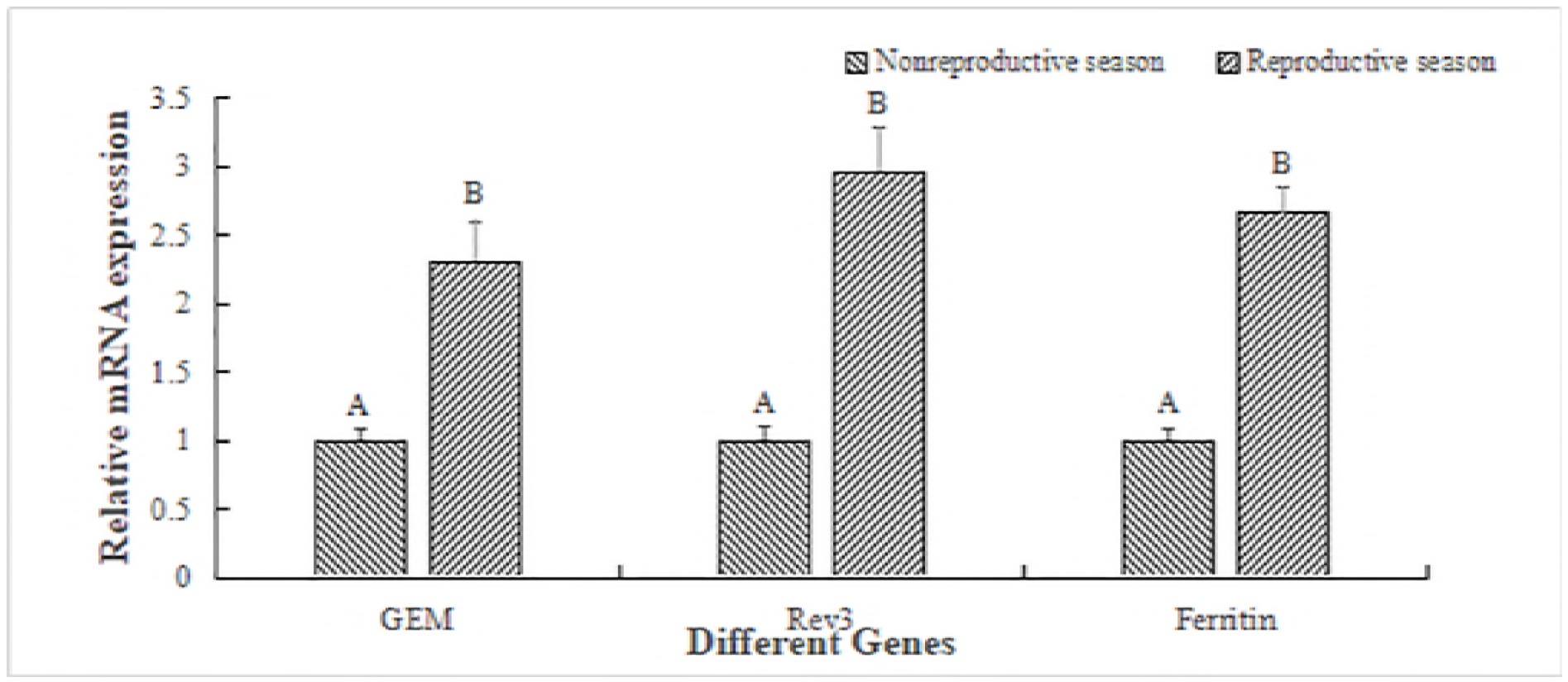
Expression characterizations at various reproductive cycle of testis were revealed by real-time quantitative PCR. The amount of mRNA was normalized to the β-actin transcript level. Data are shown as mean ± SD (standard deviation) of tissues from three separate individuals. Capital letters indicate expression difference of testes from control group.

### In situ Hybridization analysis

To analyse the functions of these three genes in depth, the mRNA locations were determined in testis, androgenic gland and hepatopancreas by *in situ* hybridization. The fixed tissue samples were subjected to hematoxylin and eosin (HE) staining as well as in situ hybridization. According to the HE staining, mature testis includes spermatid, spermatocyte and sperms, whereas sperms were the dominant cells. Androgenic gland consisted of funicular structure and androgenic gland cells. Hepatopancreas includes the lipid granules and hepatopancreas cells.

According to the *in situ* hybridization analysis, strong signals for Rev3 mRNA in mature testis were observed in spermatid, spermatocyte and sperm in mature testis (Figure 5), while strong signals for GME and ferritin peptide were only observed in spermatid, no signals were observed in spermatocyte and sperms (Figure 4, Figure 6). In androgenic gland, strong signals were observed in funicular structure surrounding the androgenic gland cells for all of these three genes, while no signal was directly observed in androgenic gland cells. Strong signals were observed in hepatopancreas cells for all of these three genes, rather than that in lipid granules (Figure 4; Figure 5; Figure 6). No signals were observed when sense RNA probe was used.

**Fig 4:**
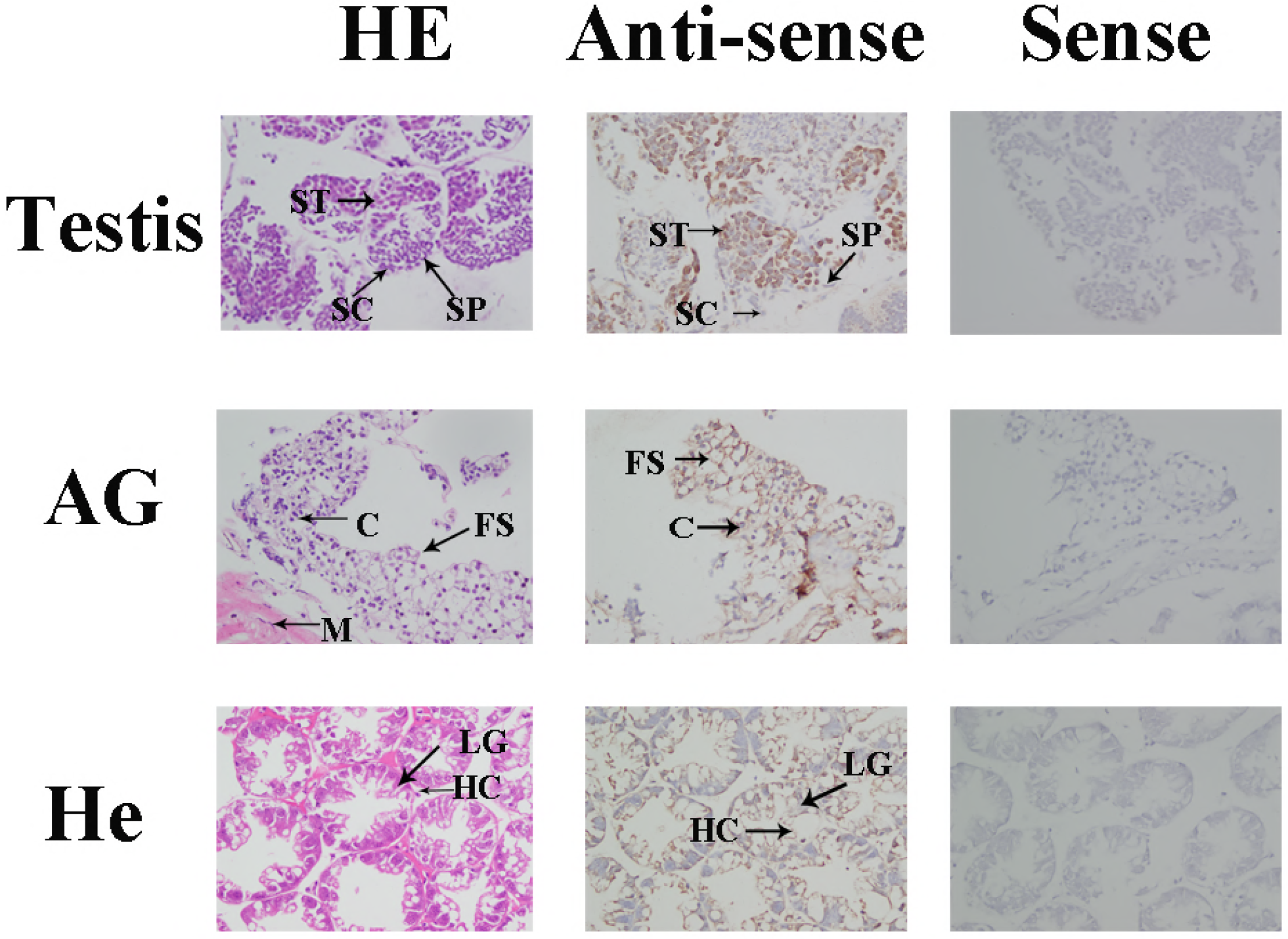
Location of GEM gene was detected in testis, androgenic gland and hepatopancreas of M. nipponense by in situ hybridization. Testis, androgenic gland and hepatopancreas were sampled at reproductive season. AG: androgenic gland; ST: spermatid; SC: spermatocyte; SP: sperm; M: muscle; C: androgenic gland cell; FS: Funicular structure; He: hepatopancreas; LG: lipid granules; HC: hepatocytes. Scale bars = 50μm.

**Fig 5:**
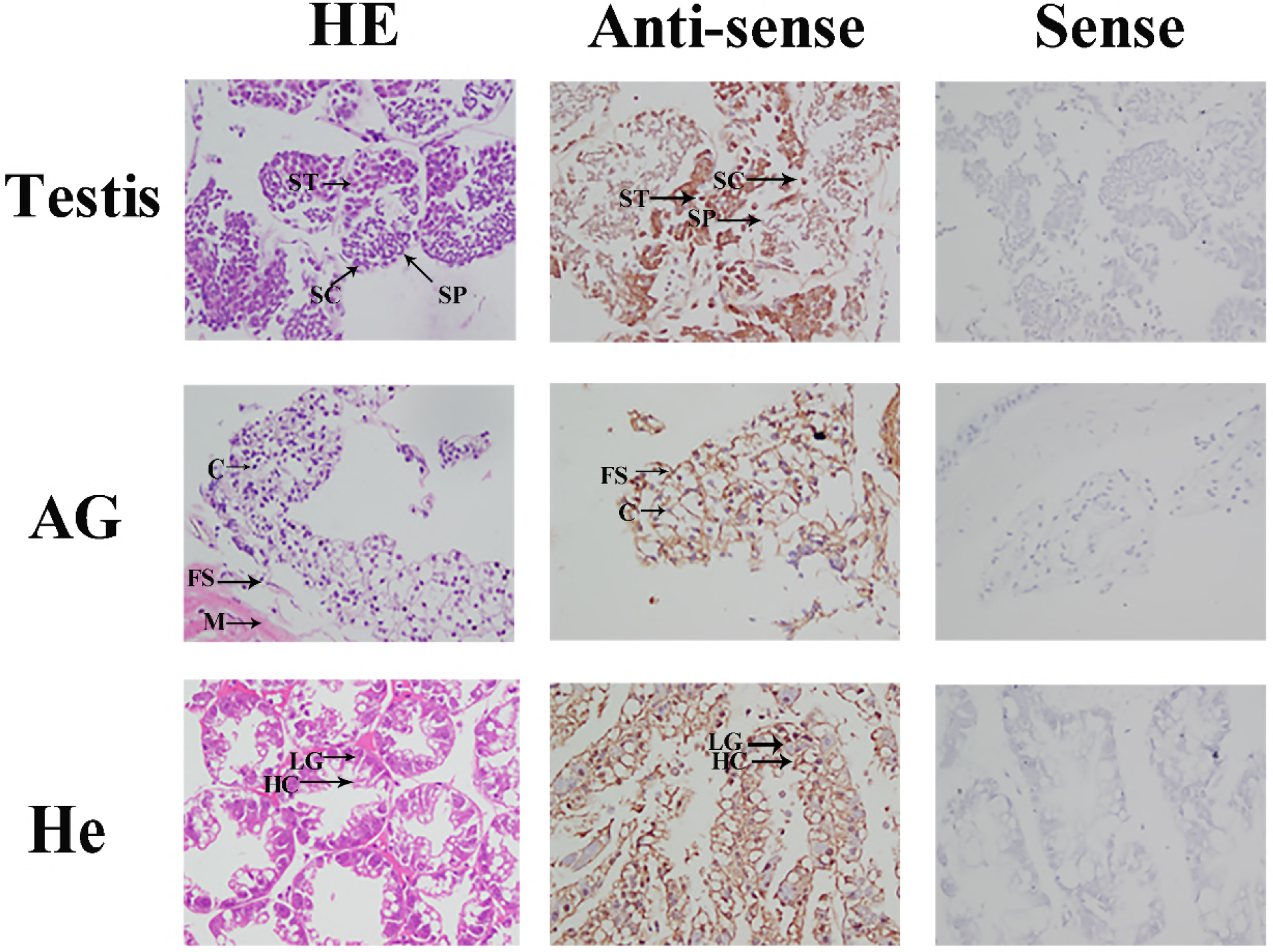
Location of Rev3 gene was detected in testis, androgenic gland and hepatopancreas of M. nipponense by in situ hybridization. Testis, androgenic gland and hepatopancreas were sampled at reproductive season. AG: androgenic gland; ST: spermatid; SC: spermatocyte; SP: sperm; M: muscle; C: androgenic gland cell; FS: Funicular structure; He: hepatopancreas; LG: lipid granules; HC: hepatocytes. Scale bars = 50 μm.

**Fig 6:**
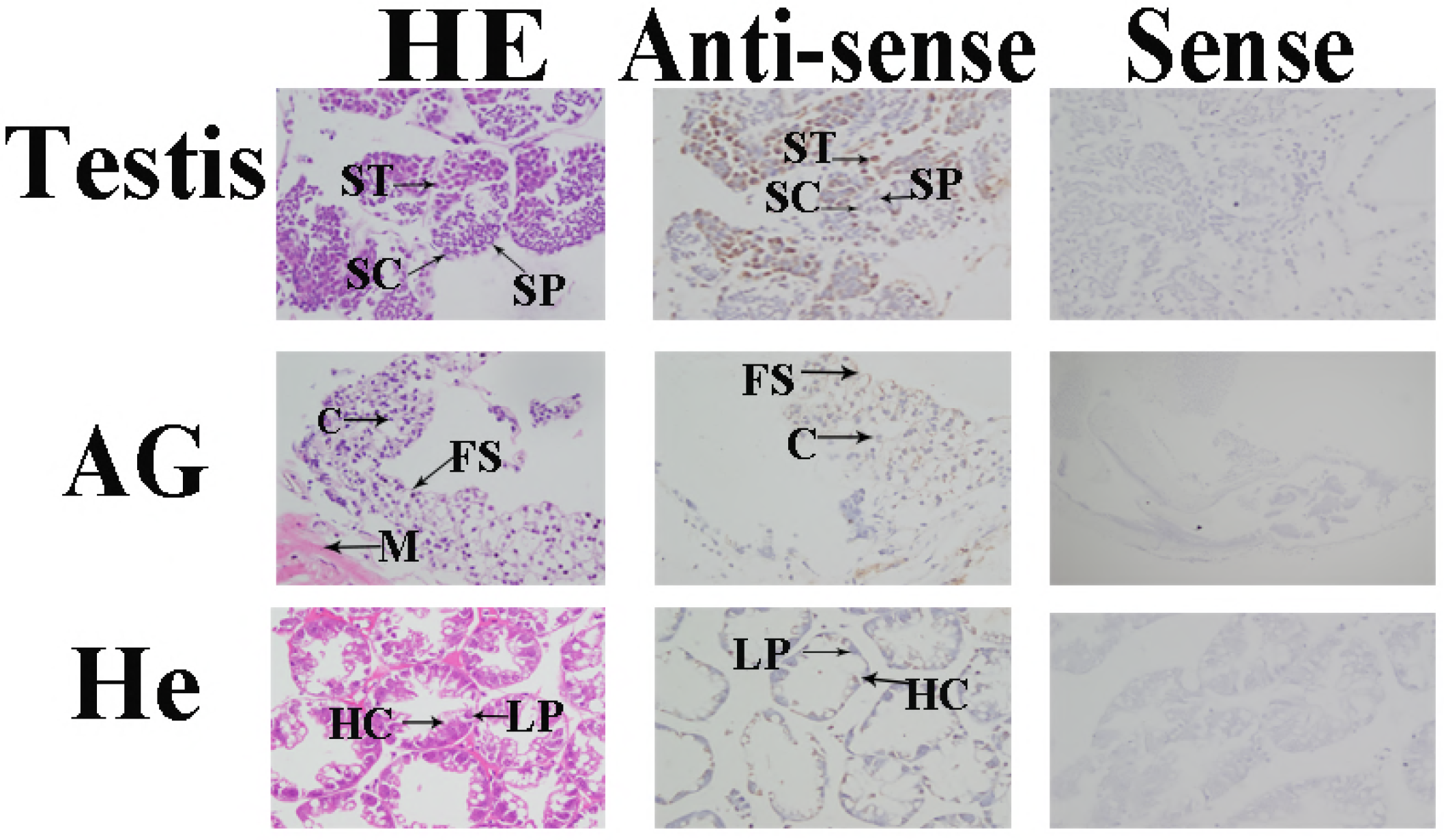
Location of ferritin peptide gene was detected in testis, androgenic gland and hepatopancreas of *M. nipponense* by in situ hybridization. Testis, androgenic gland and hepatopancreas were sampled at reproductive season. AG: androgenic gland; ST: spermatid; SC: spermatocyte; SP: sperm; M: muscle; C: androgenic gland cell; FS: Funicular structure; He: hepatopancreas; LG: lipid granules; HC: hepatocytes. Scale bars = 50 μm.

## Discussion

Three DEPs were identified from the quantitative proteomic analysis of androgenic gland from *M. nipponense* during non-reproductive and reproductive season, which showed the expressions with highest mRNA level in androgenic gland, compared with those in testis and ovary, including GEM, Rev3 and ferritin peptide. According to the important functions of androgenic gland in male sexual differentiation and development, these 3 DEPs were predicted to be strong candidate novel genes in the male sexual development in *M. nipponense.* According to the previous studies, ferritin peptide and Rev3 play essential roles in immune system maintenance [17–19]. In this study, we aimed to investigate their functions in *M. nipponense*, especially for the potentially novel roles in male sexual development. The full-length cDNA sequence of Mn-GEM was 1,018 bp with an open reading frame of 777 bp, encoding 258 amino acids. Mn-GEM has the highest identity with GEM sequence from *Acanthaster planci*, while the identity was only 41%. According to the phylogenetic analysis, Mn-GEM has dramatically long evolutionary relationship with those from other species. These suggest considerable evolutionary divergence between *M. nipponense* and other species in terms of GEM, consistent with BLASTP analysis. The partial cDNA sequence of Mn-Rev3 was 6,832 bp with an opening reading frame of 3,612 bp, encoding 1203 amino acids. The full-length cDNA sequence of Mn-Rev3 was hard to obtain by using 5’RACE cloning and homologous cloning because the absent cDNA sequence at 5’-terminal is too long and the degenerate primers were hard to designed due to long evolutionary relationship with the well-defined Rev3 sequences in other species. The full-length cDNA sequence of Mn-Rev3 will be obtained when the middle fragment is long enough. As the same as that of Mn-GEM, the similarity comparison analysis and phylogenetic analysis also showed a considerable evolutionary relationship between the Mn-Rev3 and the other well-defined sequences of Rev3 in other species. A reasonable explanation for the long evolutionary relationship with well-defined sequences is that the GEM and Rev3 sequences were only identified and isolated from limited species, and to the best of our knowledge, no previous researches related to GEM and Rev3 were found in crustacean species.

To the best of our knowledge, the functions of GEM have not been well defined and analysed. Rev3 and ferritin peptide have been proven to play essential roles in the immune system maintenance, based on the previous studies. In cultured human fibroblasts, Rev3 decrease the UV-induced mutagenesis through carrying out translesion DNA synthesis [18–19]. Ferritin peptide protect the cells from damage by excess iron through regulating the cellular and organism-wide iron homeostasis [20–23]. In addition, ferritin peptide has been also proven to play vital roles in development, cell activation, and angiogenesis [24-27]. Ferritin peptide showed the expression with highest mRNA level in hepatopancreas in *M. nipponense*, and proved to play critical roles in its innate immune defence, especially for those in cellular and organismic iron homeostasis [17]. In this study, the mRNA expression of Rev3 was the highest in hepatopancreas, and showed significant expression difference with that in other tissues (P<0.01). Rev3 also showed high expression in androgenic gland and testis. The dramatic high expression of Rev3 in hepatopancreas implies its potential roles in immune system in *M. nipponense*, which is similar with the previous studies. The ferritin peptide showed the highest mRNA expression in testis, followed by in androgenic gland and hepatopancreas. However, there is no significant expression difference between the testis, androgenic gland, and hepatopancreas. A reasonable explanation for the different from previous study is that the tissues samples may collect at different season and have individual difference. The mRNA expression of GEM was the highest in androgenic gland, followed by testis and ovary. The high expression levels of these 3 DEPs in testis and androgenic gland suggested their potentially novel functions in male sexual development of *M. nipponense*, which have not yet been identified in any species. Besides, the mRNA expressions of these 3 DEPs in reproductive seasons were dramatically higher than those in nonreproductive season, which also suggested their potential roles in testis development.

The *in situ* hybridization of ferritin peptide has been performed in several species. The *in situ* hybridization analysis in *Branchiostoma belcheri* showed that ferritin homolog is ubiquitously expressed [28]. Ferritin mRNA from *Pinctada fucata* is highly expressed at the mantle fold, revealed by the *in situ* hybridization analysis. The mantle fold is a region, playing essential roles in metal accumulation and contributing the metal incorporation into the shell [29]. In iron-loaded rats with up-regulated levels of L-ferritin mRNA, L-ferritin mRNA was localized in many organs, including colonic crypt, villus epithelial cells, small intestinal crypt and surface epithelial cells [30]. To the best of our knowledge, no previous studies focused on the *in situ* hybridization of GEM and Rev3 in any species. In this study, strong signals for Rev3 mRNA were detected in spermatid, spermatocyte and sperm in mature testis, while strong signals for GME and ferritin peptide were only detected in spermatid. These results indicated that Rev3 promotes the testis development during the whole reproductive cycle of testis, while GME and ferritin peptide only promote the activation of testis development. Strong signals were detected in the funicular structure surrounding the androgenic gland cells for all of these three genes, while no signal was directly detected in androgenic gland cells. The histological observation during the post-larval developmental stages of *M. nipponense* indicated that androgenic gland was developed at the post-larval day 10 (PL10) with the formation of funicular structure, then the androgenic gland cells were formed into the funicular structure, and the androgenic gland was matured at PL19 [31]. No signal in androgenic gland cells indicated that these three genes were not directly secreted by the androgenic gland, while the strong signals in funicular structure suggested their essential roles in the development of funicular structure, which promote and support the formation of androgenic gland cells. The strong signals in hepatopancreas cells suggested that important roles in immune system in *M. nipponense.* The similar expression pattern of these three genes in *M. nipponense* implies some relationship between each other.

## Conclusion

We cloned and characterized GEM, Rev3 and ferritin peptide from the androgenic gland of *M. nipponense*. qPCR analysis of different tissues and reproductive cycles of testis suggested that these three genes may have additional functions in male sexual development in *M. nipponense*, which is the novel functions of these three genes and has not been reported in any species yet. In situ hybridization analysis further confirmed their important roles in male sexual development in *M. nipponense*.

## Acknowledgments

We would like to acknowledge the technical support provided by Qifeng Wang and Biqiao Yao for his excellent technical support.

